# Calcium-Calmodulin Gating of Connexin43 Gap Junctions in the Absence of the pH Gating Domain

**DOI:** 10.1101/311613

**Authors:** Siyu Wei, Christian Cassara, Xianming Lin, Richard D Veenstra

**Author notes:** Corresponding Author: Richard D Veenstra, Ph.D., Department of Pharmacology, SUNY Upstate Medical University, 750 East Adams Street, Syracuse, NY 13210.

## Abstract

Intracellular protons and calcium ions are two major chemical factors that regulate connexin43 (Cx43) gap junction channels and the synergism or antagonism between pH and Ca^2+^ has been questioned for decades. In this study, we assessed whether the calcium gating mechanism occurs independently of the pH gating mechanism by utilizing the Cx43-M257 (Cx43K258stop) mutant, a carboxyl-terminal (CT) truncated version of Cx43 lacking the pH gating domain. Dual whole cell patch clamp experiments were performed on Neuroblastoma-2a (N2a) cells or neonatal mouse ventricular myocytes (NMVMs) expressing either full length Cx43 or Cx43-M257 proteins. Addition of 1 μM ionomycin to normal calcium saline reduced Cx43 or Cx43-M257 macroscopic gap junction conductance (g_j_) to zero within 15 min of perfusion, while this response was prevented by omitting 1.8 mM CaCl_2_ from the external solution or adding 100 nM calmodulin (CaM) inhibitory peptide to the internal pipette solution. The ability of connexin calmodulin binding domain (Cx CaMBD) mimetic peptides and the Gap19 peptide to inhibit the Ca^2+^/CaM gating response of Cx43 gap junctions was also examined. Internal addition of a Cx50 cytoplasmic loop CaMBD peptide (200 nM) prevented the Ca^2+^/ionomycin-induced decrease in Cx43 g_j_, while 100 μM Gap19 peptide had no effect. Lastly, the transjunctional voltage (V_j_) gating properties of NMVM Cx43-M257 gap junctions were investigated. We confirmed that the fast kinetic inactivation component was absent in Cx43-M257 gap junctions, but also observed that the previously reported facilitated recovery of g_j_ from inactivating potentials was abolished by CT truncation of Cx43. We conclude that CT pH gating domain of Cx43 contributes to the V_j_-dependent fast inactivation and facilitated recovery of Cx43 gap junctions, but the Ca^2+^/CaM-dependent gating mechanism remains intact. Sequence-specific Cx CaMBD mimetic peptides act by binding Ca^2+^/CaM non-specifically and the Cx43 mimetic Gap19 peptide has no effect on this chemical gating mechanism.

## Introduction

Formed by a 20 member protein family, gap junctions are composed of a hexamer of connexin (Cx) subunits in each cell membrane that dock extracellularly to form an intercellular aqueous pore that facilitates the passage of ions, second messengers, fluorescent dyes, small nucleic acids, etc., virtually any solute under 1 kDa in molecular mass and 14 Å in width (1,2). Gap junctions are thought to be regulated by two general gating mechanisms, a fast transjunctional voltage (V_j_) gating mechanism and a slow chemical gating mechanism (3). Chemical agents known to close gap junctions include intracellular calcium ions, intracellular pH, lipophiles, protein phosphorylation, and so on (4). Since the mid-1970s, intracellular calcium ions and pH have been known to uncouple gap junctions, though opinions varied about the relative importance of each to the uncoupling process first referred to as the “healing-over” in the heart (5-8). Though synergistic actions of cytosolic Ca^2+^ and H^+^ have been reported, opposing viewpoints on the operative role of Ca^2+^ ions or protons have persisted for decades without resolution (9-12).

Definitive evidence of a pH gating mechanism associated with connexin43 (Cx43), the connexin with the most widespread expression pattern, was provided when truncation of last 125 amino acids of the cytoplasmic carboxyl terminus (CT) abolished the pH sensitivity of Cx43 gap junction conductance (g_j_) (13). This pH gating mechanism for Cx43 gap junctions was further defined by the demonstration that the distal CT pH gating particle binds to a receptor domain located in region 119-144 of the Cx43 cytoplasmic loop (CL) in a pH-dependent manner (14). The calcium gating hypothesis evolved to the action of calmodulin (CaM) based on the inhibition of gap junction uncoupling by calmodulin (CaM) inhibitors and evidence that CaM binds to Cx32 and lens gap junctions (15-17). Since the identification of CaM binding domains (CaMBDs) on the cytoplasmic amino- and carboxyl termini of Cx32, additional connexin CaMBDs have been identified on the CL domain of Cx43, Cx50, Cx46 (sheep Cx44), and Cx45 (18-21). Previous studies in this laboratory indicated that Ca^2+^/CaM causes a gated closure of Cx43 and Cx50 gap junctions as evidenced by the reduced open probability of gap junction channels during perfusion of coupled cell pairs with the calcium ionophore ionomycin (20,22).

In this study, we tested the ability of the Ca^2+^/CaM to cause the gated closure of Cx43 gap junctions using the previously published CT-truncated version of Cx43, Cx43-M257 expressed in mouse neuro2a (N2a) cells. We also employed the Cx43^+/K258stop^ mouse, which heterologously expresses the Cx43-M257 (K258stop) protein (23), to examine this chemical gating mechanism using an endogenous expression system, homozygous Cx43-M257 neonatal mouse ventricular cardiomyocytes (NMVMs). We found that superfusion of Cx43-M257 expressing cell pairs with 1.8 mM CaCl_2_ saline containing 1 μM ionomycin inhibited gap junction conductance (g_j_) by 100% in a calcium- and calmodulin-dependent manner. We further tested the ability of connexin CaMBD mimetic peptides to inhibit the Cx-Ca^2+^/CaM gating mechanism in a sequence-specific manner and found that 200 nM of the higher CaM affinity Cx50-3 peptide was sufficient to block the Ca^2+^/CaM-induced uncoupling of Cx43 gap despite the Cx43-3 CaMBD site possessing a distinct sequence from the Cx50 CaMBD. Conversely, the Gap19 peptide, which targets a CL sequence immediately adjacent to the known Cx43 CaMBD site (24), did not inhibit the Ca^2+^/CaM-induced uncoupling process. Finally, we also examined the V_j_-gating properties of Cx43-M257 gap junctions in paired NMVMs and found that the fast kinetic component of the V_j_-dependent gating mechanism was abolished as previously reported in *Xenopus* oocytes and N2a cells (25,26). The probability of the 50 pS subconductance state of Cx43 gap junction channels was similarly reduced in favor of the 100 pS main open state of the channel.

Additionally, the increased slope g_j_ during the recovery from V_j_-dependent inactivation previously observed only in primary NMVM gap junctions was also eliminated by the CT truncation of Cx43. Our findings indicate that the Ca^2+^/CaM-dependent gating mechanism of Cx43 gap junctions remains intact after deletion of the CT pH gating domain, that sequence-specific Cx CaMBD peptides function as non-specific CaMBDs to bind CaM, that the Gap19 peptide does not influence this gating mechanism, and the Cx43 distal CT domain is somehow involved in the “facilitated” recovery of g_j_ after V_j_-dependent inactivation.

## Materials and Methods

### Cell Cultures

Murine Neuro2a neuroblastoma (N2a) cells, grown to 70-90% confluency, were cultured in 12 well culture dishes containing MEM media supplemented with 10% fetal bovine serum and transiently transfected with 1 μg of plasmid cDNA containing full-length (WT) Cx43 or Cx43-M257 cDNA sequences using Lipofectamine2000 and OptiMEM according to manufacturer’s directions (ThermoFisher Scientific). Full length Cx43 pTracer-CMV2 and Cx43-M257 pIRES2-EGFP plasmids were purified using the EndoFree plasmid minikit (Qiagen). N2a cells were lightly trypsinized after 4 hrs and plated in 35 mm culture dishes overnight for patch clamp electrophysiology studies the next day.

All cardiomyocyte experiments were performed on enzymatically dissociated neonatal C57BL/6 murine ventricular myocytes cultured for 48–72 h according to published procedures (27). The newborn mice were euthanized under isoflurane anesthesia in accordance with procedures approved by the Institutional Animal Care and Use Committee (IACUC) conforming to the Guide for the Care and Use of Laboratory Animals published by the US National Institutes of Health (NIH Publication No. 85-23, revised 1996). Newborn litters from heterozygous Cx43-M257 (Cx43K258stop) mouse matings were collected and the ventricle and tail of each newborn pup was numbered for separate enzymatic dissociation and genotyping. Genomic DNA was obtained using the DNeasy Blood and Tissue kit (Qiagen) and subjected to polymerase chain reaction (PCR) analysis using Taq polymerase (ThermoFisher Scientific) and M257 forward (5’-CAA AAC ACC CCC CAA GGA ACC TAG) and reverse (5’ GCA TCC TCT TCA AGT CTG TCT TCG) primers as originally described (23).

### Electrophysiology

Gap junctional current (I_j_) recordings were acquired using dual whole cell patch clamp procedures (28). N2a or neonatal murine ventricular myocyte (NMVM) cell pairs were held at - 40 mV and a voltage step (ΔV) was applied to one cell while maintaining the holding potential (V_h_ = -40 mV) of the partner cell. The macroscopic junctional conductance (g_j_) was calculated by the equation: g_j_ = -ΔI_2_/[(V_1_ + ΔV) - V_2_] where I_1_ and I_2_ are whole cell currents, V_1_ and V_2_ are command voltages, and (V_1_ + ΔV) - V_2_ = the transjunctional potential, V_j_. For all experiments, g_j_ was normalized by dividing the time-dependent g_j_ by the initial g_j_ measurement, or G_j_ = g_j_/g_j,initial_. To more accurately calculate g_j_, the whole cell patch electrode resistances, R_el1_ and R_el2_, were subtracted from the g_j_ = -ΔI_2_/V_j_ calculation where the actual g_j_ = apparent g_j_ – (1/R_el1_ + 1/R_el2_). R_el_ = t_cap_/C_input_ where τ_cap_ is the single cell capacitive current decay time constant and C_input_ is the input resistance of each cell. The bath saline contained (in mM): 142 NaCl, 1.3 KCl, 4 CsCl, 2 tetraethylammonium chloride, 0.8 MgSO_4_, 0.9 NaH_2_PO_4_, 1.8 CaCl_2_, 5.5 dextrose, and 10 HEPES (titrated to pH 7.4 with 1 N NaOH). 1.8 mM CaCl_2_ was omitted from the Ca^2+^-free saline. Ionomycin (1 μM, I3909, Sigma-Aldrich) was added to the perfusion saline fresh daily. The internal pipette solution (IPS) contained (in mM): 140 KCl, 1.0 MgCl_2_, 0.3 KBAPTA, 25 HEPES (titrated to pH 7.40 with 1 N KOH).

The peptide inhibition studies of Ca^2+^/CaM-dependent uncoupling of Cx43 gap junctions was performed by adding the inhibitory peptides to the IPS of both cells on the day of use at concentrations dependent upon their measured K_d_. Calmodulin (CaM) inhibition was achieved by adding 100 nM CaM inhibitory peptide corresponding to the CaMBD of CaM kinase II (Enzo Life Sciences, #BML-P200-0500) with a reported K_d_ of 52 nM. The Cx50 CaMBD peptide, corresponding to the Cx50 cytoplasmic loop sequence

141-SSKGTKKFRLEGTLLRTYVCHIIFKT-166 and its scrambled control (FKLYKCISFGGTEITTRSHVLTKKRL) were published previously (20). The CaMBD peptide was added to the IPS at a concentration of 200 nM. The Gap19 peptide (Sigma-Aldrich, #SML1426) was applied intracellularly at a concentration of 100 μM (24).

### Immunocytochemistry

For immunocytochemical studies, NMVMs were cultured on poly-l-lysine coated coverslips for 48-72 hrs and immunolabeled for Cx43 protein using previously published procedures (27). Coverslips were rinsed in phosphate-buffered saline (PBS, pH 7.0), fixed with 4% paraformaldehyde in PBS, rinsed, permeabilized with 1% Triton X100 in PBS, and blocked with 2% goat serum in 1% Triton X100 PBS, all for 15 min at room temperature. Anti-Cx43 rabbit polyclonal amino-terminal (Abgent, #AP1541b) and mouse monoclonal carboxyl-terminal (ThermoFisher #35-5000) antibodies were diluted 1:200 in 2% goat serum, 1% triton X-100 PBS and added to the 12 mm diameter coverslip overnight at 4 °C. Each well was rinsed 3-5 times with PBS the next day and incubated with goat anti-rabbit Alexa Fluor 488 (ThermoFisher #11008) and anti-mouse Alexa Fluor 546 (ThermoFisher #11003) secondary antibodies, diluted 1:1500 in 10% goat serum PBS, for 30 min at room temperature. After rinsing, the cells were labeled with DAPI for 10 min, rinsed with PBS and finally with pure water, blotted dry, and mounted on a glass slide using ProLong antifade mounting reagent and cured overnight in the dark.

Sealed glass coverslips were viewed with an Olympus IX-70 microscope using a Sutter Instruments LS 175 W Xenon arc lamp epifluorescence illumination system and Lambda 10–2 filter wheel controller with 484/15 or 555/25 nm band pass excitation filters and FITC or FITC/Cy3/Cy5 dichroic mirror/emitter filter sets (Chroma Technology Corp, #41001 or #62005). DNA staining was observed with a 365/10 nm excitation filter and DAPI 460/50 nm dichroic mirror/emitter filter set (Chroma, 31000v2-dapi-hoechst-amca). Fluorescent micrographs were acquired with an Andor iXon 885 ECCD camera using Imaging Workbench 6.0 software (INDEC Systems, Santa Clara, CA). Exported TIF files were background subtracted (≈10%) and green/red/blue color processed using ImageJ software. Magnification was 600X using an Olympus PlanApo 1.40/0.17 aperture 60X oil immersion and 10X C-mount objectives.

## Results

To examine the Ca^2+^/CaM gating of Cx43-M257 gap junctions, we employed the same ionomycin perfusion assay from our previous study on the full length wild-type (WT) Cx43 (22). When coupled WT or Cx43-M257 N2a cell pairs were superfused with normal saline containing 1 μM ionomycin and 1.8 mM CaCl_2_, junctional currents (I_j_) in response to a -20 mV V_j_ pulse declined steadily from initial levels to zero during 8-12 min of perfusion at a rate of 1 ml/min (Fig. 1A, B). The average junctional conductance (g_j_) was 26.4 ± 6.2 nS for the WT Cx43 and 29.1 ± 7.7 nS for the Cx43-M257 cell pairs (mean ± SEM, n = 7, 6). The average normalized G_j_ (= g_j(time)_/g_j,initial_) declined to 0 nS within 8 min from the onset of perfusion for both WT Cx43 and Cx43-M257 gap junctions (Fig. 1D,E). Omission of the 1.8 mM CaCl_2_ from the perfusate or the addition of 100 nM CaM (CaMKII 290-309) inhibitory peptide to the whole cell patch pipette solutions prevented the rundown of Cx43-M257 I_j_ (Figs. 1C, E, F). The average g_j_ was 18.6 ± 6.9 nS for the zero Ca^2+^ and 30.3 ± 6.9 nS for the CaM inhibitory peptide Cx43-M257 experiments (mean ± s.e.m., n = 6, 6). These results are consistent with previous findings using the full length Cx43 and are indicative of a calcium/calmodulin-dependent gating mechanism even with the absence of the Cx43 pH gating domain (13,22).

**Figure 1.**
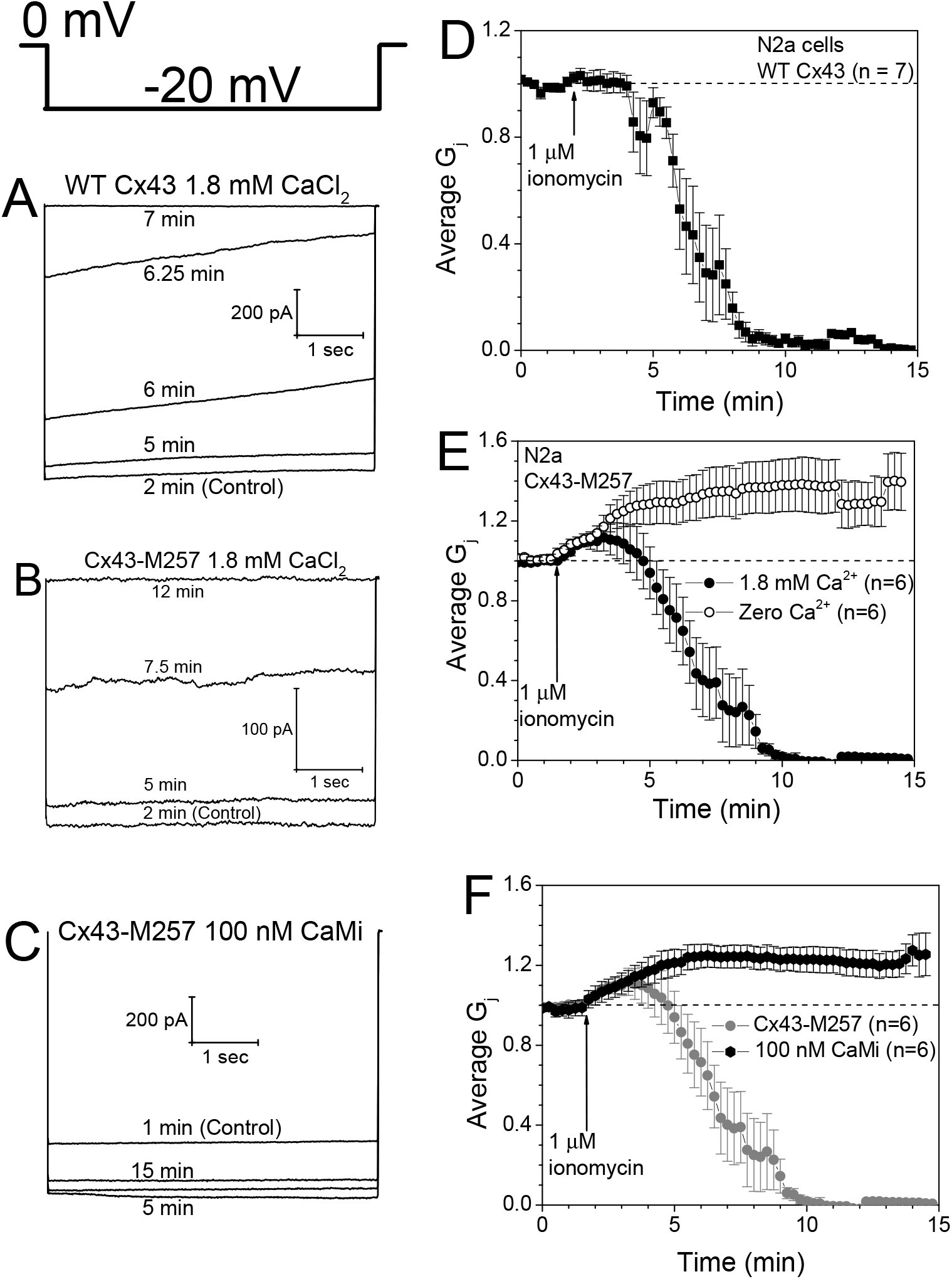
Calcium/calmodulin-dependence of Cx43 gap junctions expressed in N2a cells. **A-C**, Whole cell current traces from the partner cell 2 of an N2a cell pair during a -20 mV, 5 sec transjunctional (V_j_) voltage pulse (inset, upper left) applied to cell 1 before (control) and during perfusion with 1 μM ionomycin, 1.8 mM CaCl_2_ saline. Current traces from representative experiments illustrate the decline in junctional current (I_j_ = -ΔI_2_) during ionomycin perfusion of an N2a cell pair expressing either full-length wild-type Cx43 (**A**) or carboxyl-tail truncated Cx43 (Cx43-M257, **B**) gap junctions, or the prevention of the calcium-induced decline in I_j_ by calmodulin (CaM) inhibition (**C**). **D**, The average (mean ± SEM, n = 7) decline in junctional conductance (G_j_) during CaCl_2_/ionomycin perfusion of N2a-Cx43 cell pairs illustrating 100% G_j_ inhibition within 15 min of perfusion. **E**, Average G_j_ of N2a-Cx43-M257 cell pairs (n = 6) during perfusion with 1.8 mM CaCl_2_ saline + ionomycin (∎) illustrating complete uncoupling in the absence of the Cx43 pH gating domain, and the prevention of G_j_ inhibition by excluding CaCl_2_ from the 1 μM ionomycin saline perfusate (〇, n = 6). **F**, Inclusion of 100 nM CaM inhibitory peptide in the patch pipettes of both cells also prevents inhibition of N2a-Cx43-M257 G_j_ by calcium/ionomycin perfusion.

The above Cx43-M257 experiments were performed using the exogenous N2a cell expression system and a Cx43K258stop mouse was later developed that expresses the CT-truncated (K258stop = M257) form of Cx43 in a heterozygous manner (23). Thus, mouse ventricular cardiomyocytes were cultured from the hearts of newborn littermates from heterozygous Cx43^+/K258stop^ mice matings. The genotype of each newborn pup was determined by PCR analysis of tail DNA samples and confirmed by immunocytochemical labeling of cultured cardiomyocytes using amino-terminal and carboxyl-terminal anti-Cx43 antibodies (Fig. 2A,B). The Cx43-NT antibody recognizing both forms of Cx43 was secondarily labeled with Alexa Fluor488 (green) and the Cx43-CT antibody, which would recognize only the full-length WT Cx43, was labeled with Alexa Fluor546 (red). Homozygous Cx43-M257 cardiomyocytes were devoid of Cx43-CT immunolabeling (Fig. 2A) whereas the WT Cx43 cardiomyocytes were immunolabeled by both anti-Cx43 antibodies (Fig. 2B). Perfusion of Cx43-M257 and WT Cx43 paired cardiomyocytes with 1.8 mM CaCl_2_ + 1 μM ionomycin saline resulted in complete uncoupling within 8 min of perfusion, similar to the results obtained in N2a cells. The average g_j_ was 54.2 ± 6.2 and 39.3 ± 4.7 nS for the Cx43-M257 and WT-Cx43 cardiomyocyte pairs (n = 7, 7).

**Figure 2.**
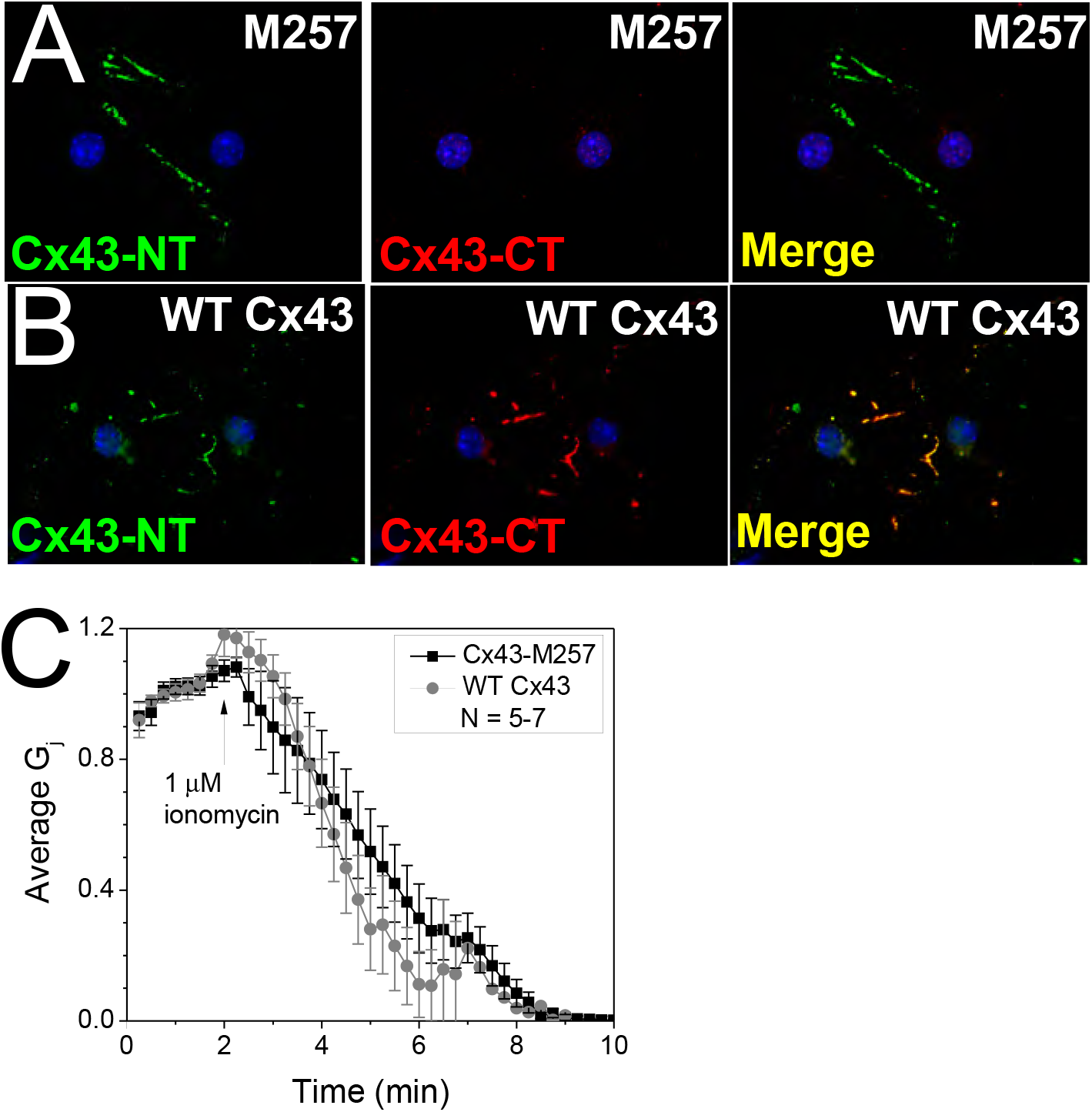
Calcium/CaM-dependent uncoupling of Cx43-M257 ventricular cardiomyocytes. **A**, Immunofluoresent labeling of Cx43-M257 cardiomyocyte gap junctions with anti-Cx43 aminoterminal (NT) and Alexa Fluor488 antibodies (green) and anti-Cx43 carboxyl-terminal (CT) and Alexa Fluor555 antibodies (red) illustrating the lack of the Cx43 CT domain in cardiomyocytes cultured from homozygous Cx43-M257 neonatal mice. **B**, Immunocytochemical labeling of wild-type Cx43 gap junctions illustrating the presence of the NT and CT domains in the littermate control cardiomyocytes. **C**, The average G_j_ of both wild-type and Cx43-M257 cardiomyocyte gap junctions was completely inhibited by calcium + ionomycin saline perfusion, confirming the results obtained by exogenous expression of Cx43 constructs in N2a cells.

We had previously shown that connexin mimetic peptides corresponding to the CaMBD of Cx43 and Cx50 were capable of preventing the uncoupling of their respective gap junctions during Ca^2+^-ionomycin perfusion (20,22). To test the specificity of these Cx CaMBD mimetic peptides, we tested different concentrations of the Cx50-3 peptide in WT Cx43 N2a cell pairs (Fig. 3A). We found that 200 nM Cx50-3 peptide was sufficient to completely prevent the decline in G_j_ during ionomycin perfusion of Cx43 gap junctions. The scrambled control Cx50-3 peptide, however, failed to prevent uncoupling during 10-12 min of Ca^2+^-ionomycin perfusion. These results suggest that any Cx mimetic peptide that is capable of binding CaM will interfere with the Ca^2+^/CaM-dependent gating mechanism in a connexin non-specific manner. The average g_j_ was 14.9 ± 5.7 and 34.4 ± 8.1 nS for the Cx50-3 and scrambled control peptide experiments (n = 5, 3).

**Figure 3.**
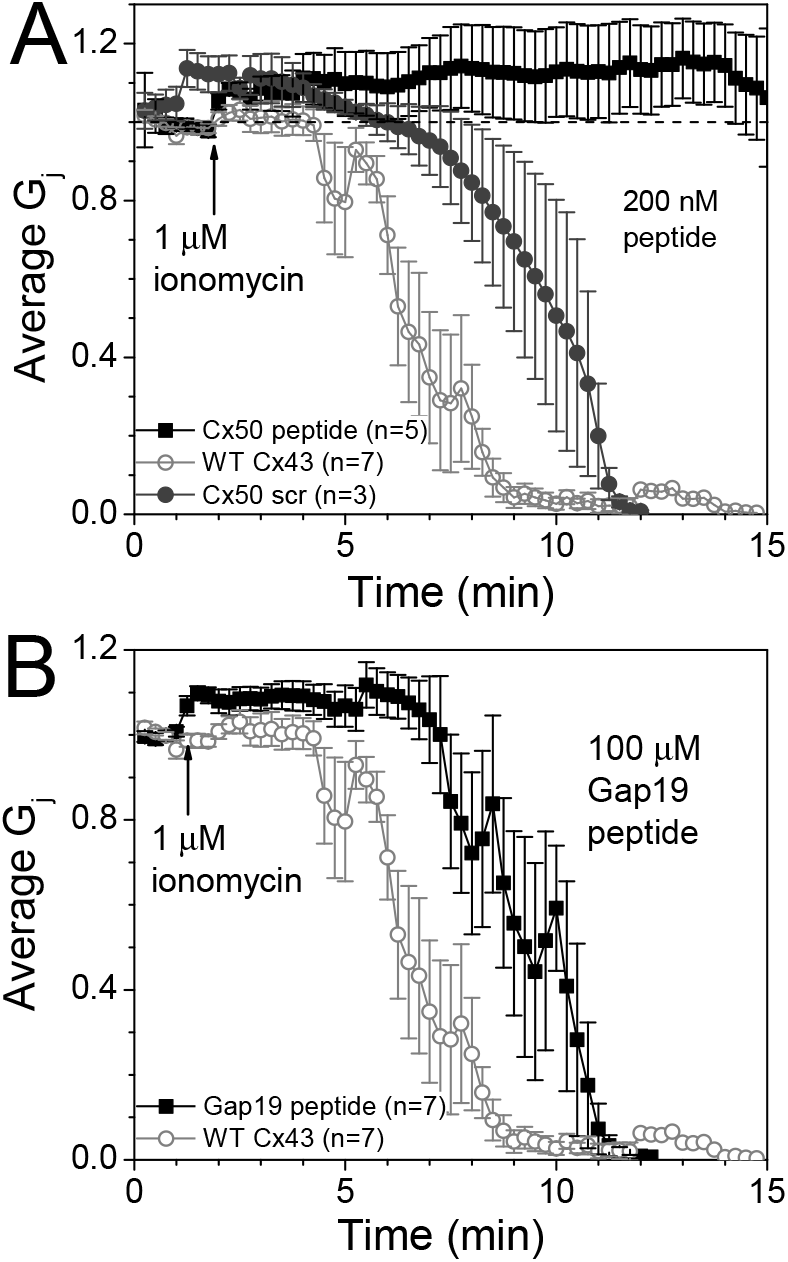
Effects of Connexin mimetic peptides on calcium/CaM-dependent uncoupling of Cx43 gap junctions. **A**, Connexin mimetic peptide of a confirmed Cx50 cytoplasmic loop (CL) calmodulin binding domain (CaMBDs) previously shown to inhibit Ca^2+^/CaM-dependent uncoupling was tested on Cx43 wild type gap junctions (20). Complete inhibition of Cx43 gap junction uncoupling was achieved with 200 nM Cx50-3 mimetic peptide added to both patch pipettes, but not the scrambled control Cx50 CaMBD peptide. **B**, A distinct Cx43 CL mimetic peptide reported to inhibit Cx43 hemichannel activity, but not Cx43 gap junction coupling, was examined for possible effects on Cx43 Ca^2+^/CaM-induced uncoupling (24). 100 μM Gap19 peptide added to both whole cell patch pipettes neither inhibited Cx43 G_j_ nor prevented the decline in G_j_ induced by 1.8 mM CaCl_2_/ionomycin saline perfusion.

The Cx43-3 CaMBD (136-158) is adjacent to the CL “L2” (119-144) region identified as the receptor for the CT pH gating domain of Cx43 (14). Recently, a Gap19 peptide targeting the central portion of the L2 region, 128-KQIEIKKFK-136, was developed and shown to inhibit Cx43 hemichannel function without inhibiting g_j_ (24). Given the close proximity of the Gap19 region to the known Cx43 CaMBD, we wanted to test for possible effects of the Gap19 peptide on the Ca^2+^/CaM-dependent uncoupling of Cx43 gap junctions. In seven Ca^2+^-ionomycin perfusion experiments, inclusion of 100 μM Gap19 peptide in both patch pipettes did not prevent the uncoupling of WT Cx43 N2a cells pairs (Fig. 3B). The average g_j_ was 31.8 ± 5.9 nS (n = 7). These results suggest that Gap19 does not affect the Ca^2+^/CaM-dependent gating mechanism of Cx43 gap junctions.

Truncation of the Cx43 CT was also reported to alter the V_j_-gating of Cx43 gap junctions by eliminating the “fast” V_j_-gating mechanism to the 50 pS subconductance state of Cx43 gap junction channels (25,26). We had previously observed a hysteresis in the steady state G_j_–V_j_ curve only in NMVMs termed “facilitation” wherein a slow (200ms/mV) V_j_ ramp from ±120 mV back to 0 mV V_j_ resulted in a higher linear slope conductance at low V_j_ values than measured initially with increasing ±V_j_ values from 0 mV (27). The previous V_j_-gating studies of Cx43-M257 gap junctions were performed in *Xenopus* oocytes and N2a cells. Thus, we examined the V_j_-gating properties of myocardial Cx43-M257 gap junctions using cultured NMVMs. Application of the slow ±120 mV V_j_ ramp revealed the loss of facilitation in Cx43^+/K258stop^ ventricular cardiomyocyte gap junctions during the return recovery phase of the G_j_–V_j_ curve (Fig. 4A, B). The average g_j_ of these V_j_-gating experiments was 7.2 ± 1.4 nS since V_j_-dependent inactivation can be detected only in low g_j_ cell pairs.

**Figure 4.**
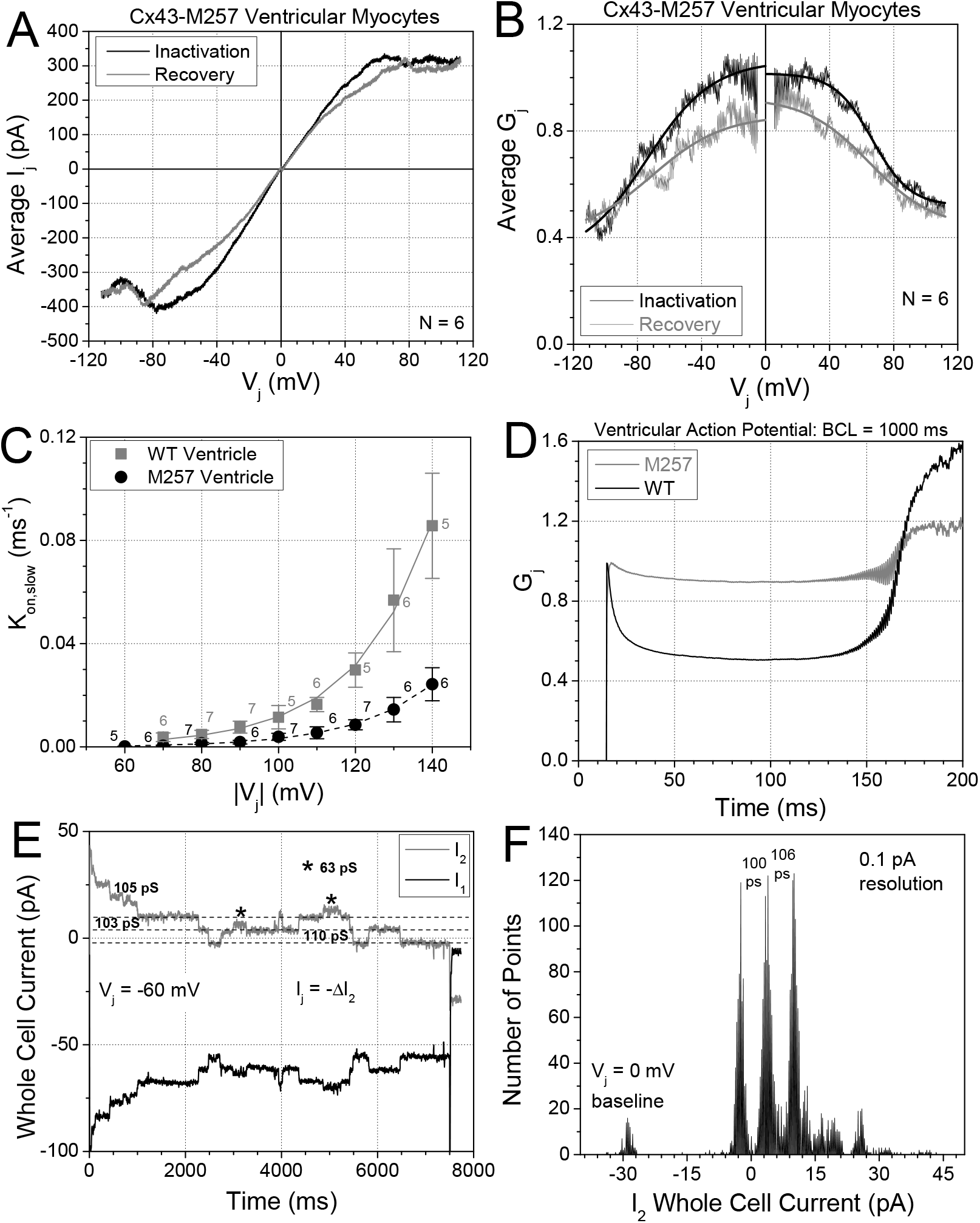
V_j_-gating and channel conductance properties of Cx43-M257 ventricular gap junctions. **A**, Average junctional current – transjunctional voltage relationship (I_j_ – V_j_) from six Cx43-M257 ventricular cardiomyocyte cell pairs obtained during a continuous 200 msec per mV V_j_ ramp from 0 to ± 120 mV (inactivation) and back to 0 mV (recovery). Absent from these traces is the increased linear slope conductance during the recovery phase observed in wild type Cx43 ventricular cardiomyocyte gap junctions (27). **B**, Corresponding average G_j_ – V_j_ inactivation (—) and recovery (—) curves depicting the typical bell-shaped curve with Boltzmann equation fits of the data (see Table 1). Again, the lack of the “facilitation” of G_j_ is apparent, wherein facilitation is defined as a peak G_j_ during the recovery phase > 1 since G_j_ is normalized to the actual junctional conductance (g_j_) value for each cell pair at V_j_ = 0 mV prior to the application 200 msec/mV V_j_ inactivation and recovery ramps. **C**, The kinetics of V_j_-dependent inactivation of wild type (WT) and Cx43-M257 ventricular cardiomyocytes were determined from 5-7 cell pairs in response to V_j_ pulses from -60 to -140 mV in 10 mV increments according to the methods of Lin et al. (27). Exponential fits of the slow inactivation rates (K_on,slow_) from WT mice revealed an e-fold change in rate for every 17.2 ± 0.9 mV compared to 22.8 ± 1.7 mV for Cx43-M257 myocytes. The fast inactivation component was absent in Cx43-M257 myocytes, consistent with previous findings (25,26). **D**, G_j_ inactivation during a 1 Hz ventricular action potential waveform from WT (—) and Cx43-M257 (—) cardiomyocytes. Owing to the slower inactivation kinetics and reduced V_j_-sensitivity of the Cx43-M257 gap junctions, V_j_-dependent inactivation of G_j_ during the action potential was reduced to 10% compared to 50% for WT Cx43 cardiomyocyte gap junctions. **E**, Gap junction channel activity observed in a Cx43-M257 myocyte cell pair during one 7.5 sec, -60 mV V_j_ pulse. Larger unitary current events corresponded to single gap junction channel conductances (γ_j_) of ≈100 pS were readily observed, but brief smaller unitary currents (*) with γ_j_ ≈ 60 pS were also observed. **F**, All points histogram of the entire I_2_ current trace shown in panel E illustrates the dominance of the 100 pS channel current events with smaller, less frequent events being lost in the noise of the dual whole cell current recording.

We also examined the kinetics of the V_j_-dependent activation in NMVM cell pairs using -V_j_ pulses and the ventricular action potential as previously described. The inactivation kinetics during V_j_ pulses from -60 to -140 mV confirmed the loss of the fast inactivation time constant and the presence of a distinct slow inactivation time constant relative to WT Cx43 NMVM gap junctions (Fig. 4C). The elimination of the Cx43 fast inactivation time constant also eliminated nearly all of the V_j_-dependent inactivation during the 1 Hz simulated ventricular action potential (Fig. 4D). The inactivation rates (K_on_) were calculated using the expression K_on_ (ms^-1^) = 1/τ_inactrvation_ = A^0^•exp(V_j_/V_k_) + C where A^0^ is the rate constant amplitude in ms^-1^, C is the minimum rate (ms^-1^), and V_k_ is the voltage constant for the inactivation rate (27,29). Exponential fits of the WT slow inactivation rates and the M257 inactivation rates yielded rates of .0000244•exp(V_j_/17.2) + 0.00236 for WT Cx43 NMVM gap junctions and 0.0000504•exp(V_j_/22.8) - 0.000058 for Cx43-M257 NMVM gap junctions. These data reveal that deletion of the CT terminus of Cx43 not only eliminated the fast inactivation of Cx43 gap junctions, but also reduced the inactivation rate and V_j_-sensitivity of the remaining slow inactivation component of Cx43 gap junctions. We also measured the unitary gap junction channel currents from three of these M257 NMVM experiments and observed an abundance of 100 pS channel events with only brief occurrences of apparent 60 pS channel events with open probabilities too low to be discernable in all points histograms of the gap junction channel current recordings (Fig. 4 E,F). Together, these data from Cx43-M257 NMVMs confirm the elimination of the fast inactivation component of Cx43 gap junctions and abundance of 100 pS gap junction channel conductance (γ_j_) events, but also quantifies the reduction in the kinetics and V_j_-sensitivity of the remaining slow inactivation component and demonstrates the removal of the of the facilitated recovery of g_j_ from inactivating potentials of Cx43 gap junctions.

## Discussion

The primary purpose of this study was to determine if the recently described Ca^2+^/CaM-dependent gating mechanism exists in the absence of the Cx43 CT pH gating domain which abolished the pH-sensitivity of Cx43 g_j_ (13,18,22). Perfusion of Cx43-M257 N2a or NMVM cell pairs with 1 μM ionomycin saline containing normal 1.8 mM CaCl_2_ routinely induced 100% inhibition of Cx43 g_j_ within 8-12 min from the onset of 1 ml/min perfusion (Figs. 1A,B,D,E and 2B). Consistent with previous findings using full length Cx43 expressed in N2a cells or wild-type NMVMs (22), omission of the 1.8 mM CaCl_2_ from the 1 μM ionomycin bath saline or addition of 100 nM CaMKII 290-309 CaM inhibitory peptide to both whole cell patch pipettes prevented the rundown of Cx43-M257 g_j_ during 13 min of perfusion (Figs. 1C,E,F). Taken together, these data support the conclusion that the Ca^2+^- and CaM-dependent chemical gating mechanism of Cx43 gap junctions does not require the presence of the Cx43 distal CT pH gating particle.

The existence of Cx CaMBDs has been demonstrated by *in vitro* CaM binding assays with sequence-specific Cx mimetic peptides comprising the entire15-26 amino acid CaMBDs of Cx32, Cx43, sheep Cx44 (Cx46), Cx45, and Cx50 (17-21). In past studies, these Cx CaMBD mimetic peptides were used as inhibitory peptides to validate the functional role of the Cx-specific sequence in the Ca^2+^/CaM gating of the parent connexin (20,22). However, like the CaMKII 290-309 inhibitory peptide which corresponds to the high affinity CaMBD of CaMKII, these peptides likely function purely on the basis of binding CaM and acting as a “CaM sponge” when added in excess to an intracellular pipette solution. To test this hypothesis, we applied increasing concentrations of the Cx50 CaMBD peptide to WT Cx43 gap junctions and found that 200 nM of the Cx50-3 peptide was sufficient to prevent the Ca^2+^/CaM-dependent inhibition of Cx43 g_j_ (Fig. 3A). This observation indicates that inhibition of the Ca^2+^/CaM gating process by these Cx-sequence specific CaMBD mimetic peptides does not necessarily prove the functionality of the corresponding domain, it also does not necessarily disprove the functional relevance of these identified Cx CaMBD domains. Additional experiments and novel approaches will be required to test the functionality of known Cx CaMBDs in a sequence-specific manner.

Another Cx43 mimetic peptide, the Gap19 nonapeptide, targeting CL residues 128-136 in the middle of the L2 pH receptor domain (residues 119-144) was shown to inhibit Cx43 hemichannel activity with an intracellular IC_50_ of 6.5 μM without affecting g_j_ at substantially higher concentrations (400 μM) presumably by interfering with the CL-CT interaction (14,24). Since the Cx43 CaMBD peptide corresponds to CL residues 136-158, it is possible that Gap19 might interfere in the Ca^2+^/CaM gating mechanism (18). To test this hypothesis, we performed the Ca^2+^-ionomycin perfusion experiments on N2a-Cx43 cell pairs with 100 μM Gap19 added to both whole cell patch pipettes (Fig. 3B). 100% inhibition of Cx43 G_j_ was still achieved within 10-12 min of perfusion and no inhibition of g_j_ was evident. Our results confirm that Gap19 does not affect Cx43 g_j_ nor the Ca^2+^/CaM-dependent uncoupling mechanism.

The V_j_-gating and γ_j_ properties of Cx43-M257 gap junctions were previously studied in exogenous *Xenopus* oocyte and N2a cell expression systems (25,26). Both studies reported a loss of the fast kinetic component of the V_j_-gating mechanism and a lower G_min_ for the steady state G_j_ – V_j_ curve. Additionally, gap junction channel recordings from Cx43-M257 N2a cell pairs revealed a loss of the low γ_j_ subconductance state, an increased open time for the remaining ≥ 100 pS main conductance state with primarily slow transitions between the open and closed states of the channel (26). Previously, we had observed a hysteresis in the steady state G_j_ -V_j_ curves obtained during the application of slow 24 sec, 0 to ±120 mV V_j_ ramps found only in primary NMVM cell pairs, not N2a-Cx43 cell pairs (27). The observed increase in the linear slope conductance at low V_j_ potentials, corresponding to the G_max_ of the steady state G_j_ – V_j_ curves, occurred during the recovery (from inactivation) phase of the ±120 mV to 0 mV V_j_ ramp in NMVMs and was termed facilitation. This facilitated recovery of G_j_ was not affected by non-specific serine/threonine protein kinase inhbitors (e.g. 12 μM H7, data not shown) nor 100 nM rotigaptide, though the inactivation phase was reduced by these treatments (29).

Thus, we examined the effect of the Cx43 CT truncation on the V_j_-gating properties on homozygous Cx43-M257 NMVMs. Our results confirmed the loss of the fast inactivation component of Cx43 gap junctions and predominance of the 100 pS main γ_j_ state of Cx43 gap junction channels seen in exogenous expression systems, but also the abolition of the facilitated recovery of G_j_ during decreasing V_j_ values after achieving steady state inactivation to G_min_ (Fig. 4A-F). We did not, however, observe a reduction in G_min_ compared to control WT NMVM gap junctions (27). The reduced rate and V_j_-sensitivity of the remaining slow kinetic inactivation component of Cx43-M257 gap junctions implies that the Cx43 CT domain contributes some of the charge to the slow V_j_ gating mechanism of Cx43 gap junctions (25). Furthermore, the deletion of the facilitated recovery of G_j_ from inactivating potentials was only previously attained by non-selective histone deacetylase inhibition (pan-HDACI) by 100 nM trichostatin A or 1 μM vorinostat implies that protein acetylation directly (e.g. Cx43 CT domain) or indirectly (e.g. tubulin?) affects the V_j_ gating properties of Cx43 gap junctions (30).

In summary, we conclude that the calcium/calmodulin-dependent chemical gating mechanism of Cx43 gap junctions does not require the distal CT pH gating particle of Cx43. The Gap19 peptide targeting a portion of the Cx43 CL pH receptor domain also does not affect the Ca^2+^/CaM gating mechanism of Cx43, but the sequence-specific Cx-CaMBD mimetic peptides function as a non-specific CaM binding domain and do not necessarily infer the modulatory function of these domains in their respective connexin-specific gap junctions. Lastly, we confirmed in Cx43-M257 cardiomyocyte gap junctions that deletion of the Cx43 CT domain eliminates the fast inactivation component and enhances the probability of the 100 pS main γ_j_ state of Cx43 gap junctions, but also eliminates the facilitated recovery of g_j_ from inactivation.

**Table 1.**
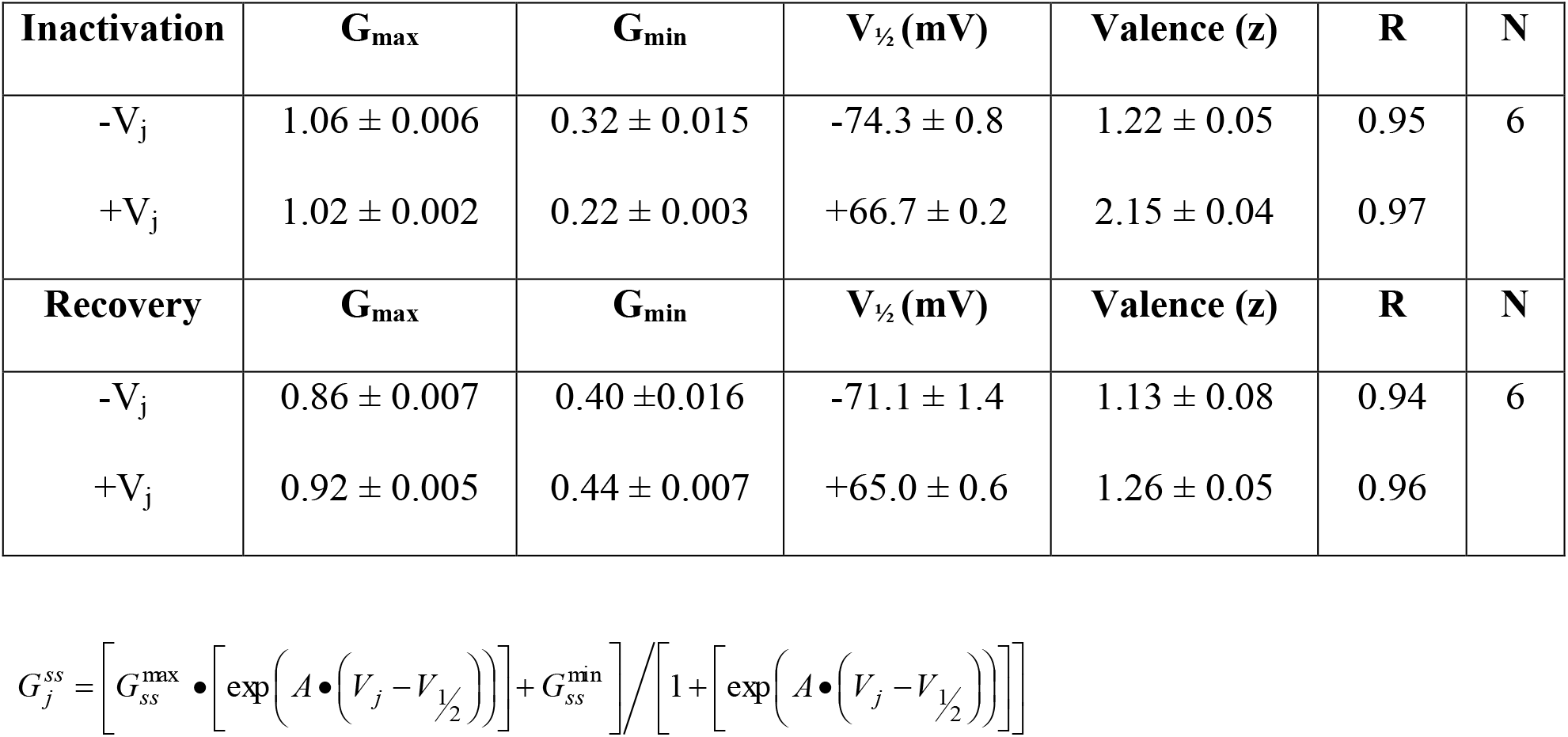

| **Inactivation** | **G<sub>max</sub>** | **G<sub>min</sub>** | **V<sub>1/2</sub> (mV)** | **Valence (z)** | **R** | **N** |
| --- | --- | --- | --- | --- | --- | --- |
| -V<sub>j</sub> | 1.06 ± 0.006 | 0.32 ± 0.015 | -74.3 ± 0.8 | 1.22 ± 0.05 | 0.95 | 6 |
| +V<sub>j</sub> | 1.02 ± 0.002 | 0.22 ± 0.003 | +66.7 ± 0.2 | 2.15 ± 0.04 | 0.97 |  |
| **Recovery** | **G<sub>max</sub>** | **G<sub>min</sub>** | **V<sub>1/2</sub> (mV)** | **Valence (z)** | **R** | **N** |
| -V<sub>j</sub> | 0.86 ± 0.007 | 0.40 ± 0.016 | -71.1 ± 1.4 | 1.13 ± 0.08 | 0.94 | 6 |
| +V<sub>j</sub> | 0.92 ± 0.005 | 0.44 ± 0.007 | +65.0 ± 0.6 | 1.26 ± 0.05 | 0.96 |  |

## Acknowledgements

We thank Dr. Steven Taffet for the gift of the Cx43-M257 pIRES2-EGFP plasmid.

We thank Dr. Karen Maass for providing us with the Cx43^+/K258stop^ mice and the methodology for genotyping these mice.

R.D.V. was supported by grants from the NIH HL042220, AHA 17GRNT33710031, Hendricks Fund, and Joseph C Georg Fund from the CNY Community Foundation.

## References

1. Harris, A. L. 2007. Connexin channel permeability to cytoplasmic molecules. Prog. Biophys. Molec. Biol. 94:120–143.

2. Valiunas, V., I. S. Cohen, and P. R. Brink. 2018. Defining the factors that affect solute permeation of gap junction channels Biochim. Biophys. Acta 1860:96–101.

3. Bukauskas, F. F., and V. K. Verselis. 2004. Gap junction channel gating. Biochim. Biophys. Acta 1662:42–60.

4. Harris, A. L. 2004. Emerging issues of connexin channels: biophysics fills the gap. Q. Rev. Biophys. 34:325–472.

5. Rose, B., and W. R. Loewenstein. 1975. Permeability of cell junction depends on local cytoplasmic calcium activity. Nature 254:250–252.

6. Turin, L., and A. Warner. 1977. Carbon dioxide reversibly abolishes ionic communication between cells of early amphibian embryo. Nature 270:56–57.

7. De Mello, W. C., G. E. Motta, and M. Chapeau. 1969. A study on the healing-over of myocardial cells of toads. Circ. Res. 24:475–487.

8. Peracchia, C. 2004. Chemical gating of gap junction channels; roles of calcium, pH and calmodulin. Biochim. Biophys. Acta 1662:61–80.

9. Spray, D. C., A. L. Harris, and M. V. Bennett. 1981. Gap junctional conductance is a simple and sensitive function of intracellular pH. Science 211:712–715.

10. Burt, J. M. 1987. Block of intercellular communication: interaction of intracellular H^+^ and Ca^2+^. Am. J. Physiol. Cell Physiol. 253:C607–C612.

11. Peracchia, C. 1990. Increase in gap junction resistance with acidification in crayfish septate axons is closely related to changes in intracellular calcium but not hydrogen ion concentration. J. Membr. Biol. 113:75–92.

12. White, R. L., J. E. Doeller, V. K. Verselis, and B. A. Wittenberg. 1990. Gap junctional conductance between pairs of ventricular myocytes is modulated synergistically by H^+^ and Ca++. J. Gen. Physiol. 95:1061–1075.

13. Morley, G. E., S. M. Taffet, and M. Delmar M. 1996. Intramolecular interactions mediate pH regulation of connexin43 channels. Biophys. J. 70:1294–1302.

14. Duffy, H. S., P. L. Sorgen, M. E. Girvin, P. O’Donnell, W. Coombs, S. M. Taffet, M. Delmar, and D. C. Spray DC. 2002. pH-dependent intramolecular binding and structure involving Cx43 cytoplasmic domains. J. Biol. Chem. 277:36706–36714.

15. Welsh, M. J., J. C. Aster, M. Ireland, J. Alcala, and H. Maisel. 1982. Calmodulin binds to chick lens gap junction protein in a calcium-independent manner. Science 216:642–644.

16. Peracchia, C., X. Wang, L. Li, and L. L. Peracchia. 1996. Inhibition of calmodulin expression prevents low-pH-induced gap junction uncoupling in Xenopus oocytes. Pflügers Arch. 431:379–387.

17. Török, K., K. Stauffer, and W. H. Evans. 1997. Connexin 32 of gap junctions contains two cytoplasmic calmodulin-binding domains. Biochem. J. 326:479–483.

18. Zhou, Y., W. Yang, M. M. Lurtz, Y. Ye, Y. Huang, H. W. Lee, Y. Chen, C. F. Louis, and J. J. Yang. 2007. Identification of the calmodulin binding domain of connexin 43. J. Biol. Chem. 282:35005–35017.

19. Zhou, Y., W. Yang, M. M. Lurtz, Y. Chen, J. Jiang, Y. Huang, C. F. Louis, and J. J. Yang. 2009. Calmodulin mediates the Ca^2+^-dependent regulation of Cx44 gap junctions. Biophys. J. 96:2832–2848.

20. Chen, Y., Y. Zhou, X. Lin, H. C. Wong, Q. Xu, J. Jiang, S. Wang, M. M. Lurtz, C. F. Louis, R. D. Veenstra, and J. J. Yang. 2011. Molecular interaction and functional regulation of connexin50 gap junctions by calmodulin. Biochem. J. 435:711–722.

21. Zou, J., M. Salarian, Y. Chen, Y. Zhuo, N. E. Brown, J. R. Hepler, and J. J. Yang. 2017. Direct visualization of interaction between calmodulin and connexin45. Biochem. J. 474:4035–4051.

22. Xu, Q., R. F. Kopp, Y. Chen, J. J. Yang, M. W. Roe, and R. D. Veenstra. 2012. Gating of connexin 43 gap junctions by a cytoplasmic loop calmodulin binding domain. Am. J. Physiol. Cell. Physiol. 302:C1548–C1556.

23. Maass, K., A. Ghanem, I. S. Kim, M. Saathoff, S. Urschel, G. Kirfel, R. Grümmer, M. Kretz, T. Lewalter, K. Tiemann, E. Winterhager, V. Herzog, and K. Willecke. 2004. Defective epidermal barrier in neonatal mice lacking the C-terminal region of connexin43. Mol. Biol. Cell 15:4597–4608.

24. Wang, N., E. De Vuyst, R. Ponsaerts, K. Boengler, N. Palacios-Prado, J. Wauman, C. P. Lai, M. De Bock, E. Decrock, M. Bol, M. Vinken, V. Rogiers, J. Tavernier, W. H. Evans, C. C. Naus, F. F. Bukauskas, K. R. Sipido, G. Heusch, R. Schulz, G. Bultynck, and L. Leybaert. 2013. Selective inhibition of Cx43 hemichannels by Gap19 and its impact on myocardial ischemia/reperfusion injury. Basic Res. Cardiol. 108:309.

25. Revilla, A., C. Castro, and L. C. Barrio. 1999. Molecular dissection of transjunctional voltage dependence in the connexin-32 and connexin-43 junctions. Biophys. J. 77:1374–1383.

26. Moreno, A. P., M. Chanson, J. Anumonwo, I. Scerri, H. Gu, S. M. Taffet, and M. Delmar. 2002. Role of the carboxyl terminal of connexin43 in transjunctional fast voltage gating. Circ. Res. 90:450–457.

27. Lin, X., J. Gemel, E. C. Beyer, and R. D. Veenstra. 2005. Dynamic model for ventricular junctional conductance during the cardiac action potential. Am. J. Physiol. Heart Circ. Physiol. 288:H1113–H1123.

28. Veenstra, R. D. 2016. Establishment of the Dual Whole Cell Recording Patch Clamp Configuration for the Measurement of Gap Junction Conductance. Methods Mol. Biol. 1437:213–231.

29. Lin, X., C. Zemlin, J. K. Hennan, J. S. Petersen, and R. D. Veenstra. 2008. Enhancement of ventricular gap junction coupling by rotigaptide. Cardiovasc. Res. 79:416–426.

30. Xu, Q., X. Lin, L. Andrews, D. Patel, P. D. Lampe, and R. D. Veenstra. 2013. Histone deacetylase inhibition reduces cardiac connexin43 expression and gap junction communication. Front. Pharmacol. 4:44.

